# HRT Atlas v1.1 database: redefining human and mouse housekeeping genes and candidate reference transcripts by mining massive RNA-seq datasets

**DOI:** 10.1101/787150

**Authors:** Bidossessi Wilfried Hounkpe, Francine Chenou, Franciele Lima, Erich Vinicius de Paula

## Abstract

Housekeeping (HK) genes are constitutively expressed genes that are required for the maintenance of basic cellular functions. Despite their importance in the calibration of gene expression, as well as the understanding of many genomic and evolutionary features, important discrepancies have been observed in studies that previously identified these genes. Here, we present Housekeeping Transcript Atlas (HRT Atlas v1.0, www.housekeeping.unicamp.br) a web-based database which addresses some of the previously observed limitations in the identification of these genes, and offers a more accurate database of human and mouse HK genes and transcripts. The database was generated by mining massive human and mouse RNA-seq data sets, including 12,482 and 507 high-quality RNA-seq samples from 82 human non-disease tissues/cells and 15 healthy tissues/cells of C57BL/6 wild type mouse, respectively. User can visualize the expression and download lists of 2,158 human HK transcripts from 2,176 HK genes and 3,024 mouse HK transcripts from 3,277 mouse HK genes. HRT Atlas also offers the most stable and suitable tissue selective candidate reference transcripts for normalization of qPCR experiments. Specific primers and predicted modifiers of gene expression for some of these HK transcripts are also proposed. HRT Atlas has also been integrated with regulatory elements from Epiregio server. All of these resources can be accessed and downloaded from any computer or small device web browsers.

## Introduction

Housekeeping (HK) genes have been classically defined as genes that are required for the maintenance of basic cellular functions, important for the existence of any cell type. Hence, they are expected to be constitutively expressed in all cell types of the organism in normal physiological condition regardless of specific cell function, cell cycle step or developmental stage (1, 2). Due to these characteristics, HK genes are useful as references of gene expression in molecular biology and computational experiments (3–9), as well as in our understanding of various structural and functional genomics and evolutionary features (10–13). In biomedical research, the importance of the precise identification of HK genes stems from the fact that these genes are used as internal controls for the calibration of quantitative PCR (qPCR), a workhorse technique in molecular biology and biotechnology laboratories used to quantitatively estimate the expression of any gene of interest under different experimental conditions (9, 14). However, the fact that HK genes are used as calibrators for these analyses implicates that the accuracy of qPCR results can be severely jeopardized if an inadequate HK gene is used. Accordingly, the selection of HK genes can well be considered one of the factors associated with the reproducibility crisis that affect biomedical science.

In the early days of qPCR, only a limited number of genes such as *GAPDH, B2M, HPRT*, actins, tubulins or rRNA genes were considered HK genes, based on their functions associated with cell maintenance, and on the observation that they tended to be constitutively expressed (15–17). Since then, these genes have been extensively used as reference genes for qPCR normalization (18–20). However, more recently, the expression of some of these genes have been shown to vary considerably across cell types and conditions, suggesting that their widespread use might not be the most accurate strategy to calibrate qPCR results (21–23). The advent of high-throughput transcriptomic technologies allowed the identification of larger lists, encompassing hundreds to thousands of putative HK genes (1, 15, 24–27). However, despite representing an improvement in the identification of HK genes, the concordance between these studies is still low (1, 15, 24–26), and false positives remain a problem (2).

Problems with these currently available HK gene lists (1, 15, 24–26) could be associated with limitations in studies that generated them, with the first being the very definition of what a HK gene is. HK genes are normally defined according to two definitions. The first one assumes that HK genes are constitutively expressed in every tissue, with a level above an arbitrary cutoff level, used to distinguish candidate HK genes from noise and/or from weakly expressed parts of the genome (2, 15, 25). The second definition emphasizes that a HK gene should present a constant and stable expression across all tissue, instead of using a universal cutoff of expression level (1, 2). However, as genes can be expressed at different level in different tissues, the former definition excludes genes that are stably expressed at different levels in different tissues. Even though the second definition extends the first one, both failed to consider alternative splicing, a fundamental aspect of transcriptome complexity. As genes may have one or more isoforms variably expressed across tissues, instead of the “one gene, one polypeptide” concept, it is possible that one gene stably expresses one transcript in a set of tissues or cells, and another transcript in other sets of tissues (1, 28, 29). Therefore, we propose that a refined definition of HK genes should be formulated as: a single constitutive gene that expresses at least one of its protein-coding transcripts at a non-zero expression level, with low variability, which may either, be constitutively expressed in all or in a subset of tissues. In the case that any of these transcripts are constitutively expressed across all tissues, they can be referred to as a HK transcript. Following the standardization of qPCR terminology provided by the minimum information for publication of quantitative real-time PCR experiments (MIQE) guidelines (14), genes used for normalization should be referred to as reference genes, not as HK genes. Accordingly, we use the term “reference transcript” to define transcripts that are suitable for qPCR normalization (i.e., fulfill the criteria of HK transcript and are stably expressed in the specific experimental condition or tissue of interest). As such, we described in this study both HK transcripts/genes for general purpose and reference transcripts only for qPCR expression normalization in a tissue-selective context. We excluded non-coding RNAs as evidences showed that they can display higher natural sample-to-sample expression variation than protein-coding genes (30), which seems to be multi-factorial. As such, it will be difficult to make prediction using these non-coding transcripts.

A second limitation of current HK gene lists refers to the fact that they were identified in studies involving a relatively low diversity of tissues and cell types, as well as low sampling, which can introduce false positives (1, 15, 24–26). And finally, technical biases from first generation microarray and Expressed Sequence Tag (EST) sequencing data could have also affected the accuracy of HK gene identification in previous studies. Briefly, besides the problem of hybridization and cross-hybridization artifacts, microarrays are also affected by dye-based detection issues and the capability to detect low abundant genes that can be physiologically relevant. Furthermore, the coverage of all possible genes by first-generation platforms is also very limited (15, 27). EST sequencing data are mainly affected by biases of cDNA libraries cloning processes which determine what transcript sequences are represented (31). In contrast to these technologies, RNA-seq technology overcomes microarray and EST limitation and provides a more comprehensive way to measure transcriptome by ultra-high-throughput sequencing (32). Together these limitations may explain the large discrepancy of currently available HK gene lists, and justify efforts to refine their definition and identification.

To fill this gap we present HRT Atlas v1.0, a web-based tool that provides access to a reliable database of human and mouse HK genes and transcripts. By combining high-quality gene expression data generated with RNA-seq from a large diversity of tissues deposited at two large public databases (GTEx and ARCHS4) (33), with a stringent detection strategy based on a refined and strict definition of what a HK gene is, HRT Atlas v1.0 offers to the research community a valuable and accurate tool to the identification of these genes and transcripts for use as calibrators of qPCR experiments, as well as for other research questions.

## Data collection

The workflow for the generation of our database (Figure 1) was based on data generated in the GTEx project (version 7) and ARCHS4 (33). These projects provide expression data from RNA-seq in a useful processed format that enables the reuse of their datasets. For the identification of human HK genes, non-normalized transcript level read counts and meta-data from 11,281 samples, including 52 non-disease sites tissues were downloaded from GTEx portal. These data were also used to identify the most reliable candidate reference genes for each cell and tissue. In addition, other 1,201 samples from 30 human tissues and cell types were obtained from ARCHS4 to increase the candidate reference genes database following a rigorous filtering strategy: (i) sequencing depth equal or higher than 20,000,000 reads; (ii) alignment rate provided by ARCHS4 higher than 70%; (iii) library generated from mRNA enrichment protocol; (iv) and construction of paired end read library. These strategies were designed to enable an accurate detection of transcript isoforms. In fact, relevant studies library have already pointed to the importance of RNA-seq depth and paired-end read construction in the estimation of alternative spliced isoforms expression level (34–36). For the identification of mouse HK genes, 507 high-quality RNA-seq data sets from 15 tissues and cells types from wild type healthy C57BL/6 control mice were manually curated from ARCHS4 following the same criteria described above.

**Figure 1:**
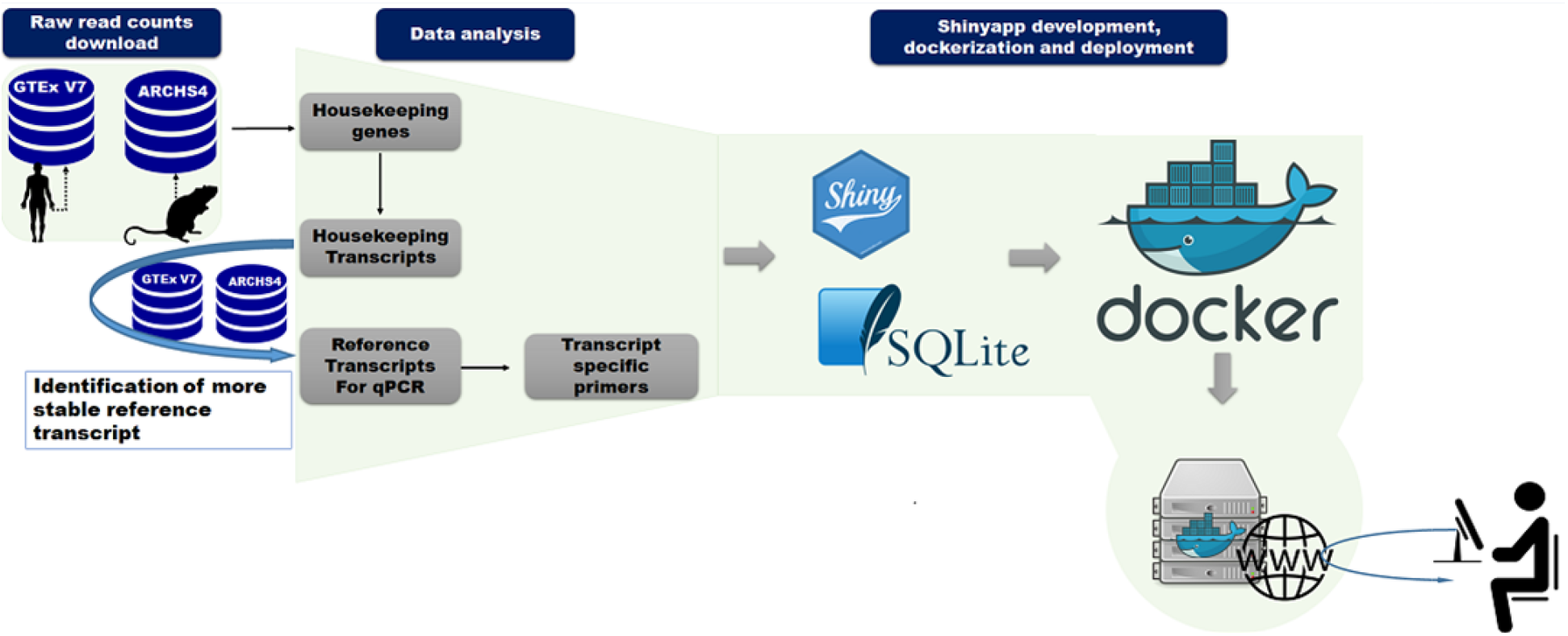
Housekeeping Transcript Atlas v1.0 workflow. The workflow of database generation and web-tool implementation are shown. After downloading and identification of HK genes using a specific algorithm, transcripts of genes with unknown pseudogenes were used to select suitable candidate reference transcripts. Some candidate reference transcript-specific primers were designed. The web-based tool was developed using Shiny package and encapsulated into docker image for deployment.

## Data processing

### Identification of HK genes

After downloading of the datasets, read counts were library-size normalized for each cell and tissue type (using TMM (Trimmed Mean of M-values) normalization factor), and the TMM normalized RPKM (reads per kilobase million) were calculated using edgeR package in R environment (37, 38). The identification of HK genes was based on the previously mentioned definition of a HK gene: specifically, to be considered a HK gene, a single gene must have fulfilled the following criteria: (i) the gene must be expressed at non-zero level in all tissue and cell types included in the analysis (that is, at least one of its protein-coding transcripts must have an expression level higher than 1 RPKM); (ii) the variability of transcript expression should be low within all tissues and cell types, as evidence by a standard deviation of the log2 RPKM < 1; (iii) the maximum fold change (MFC), represented by the ratio between maximum and average log2 RPKM of the transcript ((maximum log2 RPKM)/(average log2 RPKM)), must be lower than 2. Only transcripts with well-supported transcript models were included in the database (transcripts with Refseq and/or the Consensus Coding Sequence (CCDS) project annotation). Finally, transcripts belonging to these HK genes were considered HK transcripts if they were, in addition, constitutively expressed in all of the analyzed tissues and cells types that were analyzed.

### Identification of candidate reference transcripts

The conventional strategy of qPCR normalization used the expression of reference genes to calibrate the expression levels of genes of interest. Here we observed that only a few transcripts per HK gene fulfilled our criteria and were defined as HK transcripts. Thus, a more accurate strategy would be the normalization of qPCR experiments using reference transcripts instead of the currently used “gene model”. In order to test this hypothesis, we summarized the transcript RPKM into gene model as previously described (39). Then, genes RPKM were further submitted to our HK genes identification criteria. Overall, 1,003 genes passed our criteria, of which only 823 were common with HRT Atlas v1.0 HK genes list (Table S1). This reinforces the adequacy of our suggested normalization strategy. Further qPCR experiments are being performed to demonstrate the reliability of the transcript-based method for determining candidate reference genes.

So, to refine the list of transcripts that can be further validated as reference transcripts in qPCR experiments, reliable transcripts were selected for each tissue. Briefly, the list of identified HK transcripts was used to provide suitable reference transcripts for selected cells and tissues, for qPCR experiment normalization. Candidate reference transcripts had presented a standard deviation of the log_2_ RPKM < 0.5 and mean of RPKM greater or equal to 30. This threshold has been arbitrarily fixed to minimize the selection of transcripts which could exhibit very high or undetected Ct in a particular qPCR experimental condition. This threshold have been set as default but alternatively, user can disable the filtering option to display the full list of candidate reference transcripts, including those with mean RPKM less than 30. Finally, to ensure reliability of qPCR experiments, isoforms of genes with known pseudogenes were also excluded from the reference transcripts lists (40, 41).

## Candidate reference transcript ranking system: Score Product

Following candidate reference transcript identification, an algorithm based on the determination of a “Score product” was developed to rank these transcripts in each tissue or cell type. The algorithm was built to prioritize the most reliable transcripts for qPCR normalization. Candidate reference transcript stability was ranked based on mean RPKM, standard deviation of log_2_ RPKM and MFC value. Giving *n* transcripts, three intermediate scores (*k*=3) with equal weight were generated (score-1, score-2, and score-3 for RKPM, standard deviation and MFC, respectively for each transcript). Score-1 assigns *n* to the highest expressed transcript, estimated by mean RPKM, and 1 to the lowest expressed. Similarly, for score-2 and score-3 the highest scoring transcripts are the ones that have the lowest standard deviation or the lowest MFC. Given *sc*_*t,i*_ the score of the transcript t for the *i*-th rank, we express the Score Product (SP) via the geometric mean:

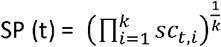

The highest scoring transcript is one that has the highest SP and is assigned to the first position (rank = 1). That way, the *n*-th transcript is the one that has the lowest SP. The Score Product is proportional to the variability.

## Primer design

In addition to the identification and ranking of candidate reference transcripts, we also manually designed specific primer pairs that can be used for qPCR normalization. These candidate reference transcript-specific primer pairs were designed to facilitate the reference transcript-based normalization strategy that we are suggesting. Primer-pairs were designed to target amplicons with length spanning between 75 and 200bp, and templates with long repeats of single bases (>4) were avoided. Each primer of a primer pair was quality checked with the following criteria: (i) GC content must be between 50 and 60%; (ii) the melting temperature (Tm) of all primer pairs must be between 50°C and 65°C with a Tm difference between primer pairs ideally lower than 3 °C; and (iii) secondary structure and primer-dimer formation were avoided. At least one of the primer-pairs was designed to span two exon junctions to minimize genomic DNA contamination, and their transcript level specificity was checked using In-Silico PCR web tool (42). Furthermore, salt concentration of 50mM (Na^+^) and divalent ion concentration of 3 mM (Mg^++^) have been used. Two annealing oligo concentrations (200 and 300nM) have been tested and 200nM was chosen for having greater or equal performance during validation. Finally, we excluded all genes with known pseudogene from the candidate reference transcripts detection pipeline. These settings didn’t be considered to completely avoid genomic DNA amplification; as such they can’t replace a rigorous pre-processing such as the use of DNase.

## Integration of HRT Atlas with predicted gene expression modifiers

In order to further guide the choice of suitable candidate reference genes for qPCR normalization, our list of human HK genes was integrated with the following resources extracted from the Harmonizome platform (43), which were downloaded and manually curated: *GEO Signatures of Differentially Expressed Genes for Diseases, GEO Signatures of Differentially Expressed Genes for Small Molecules and Connectivity Map Signatures of Differentially Expressed Genes for Small Molecules*. These datasets were used to provide users with a list of small molecules and diseases which have been previously shown to modify the expression of each HK gene, when available.

## Implementation and data access

All of the generated data were loaded into SQLite database. HRT Atlas v1.0 web tool was developed using Shiny package (44). The application front-end was implemented using HTML, CSS and Bootstrap 4. All of the components and a Shiny Server were encapsulated in a deployable Docker container running on a Unix-based operating system with Apache HTTP server. User instructions communicated to the server were processed by R and the requests are sent to the SQLite database mounted as the container volume. Figure 1 schematically represents the general workflow of the resource (Figure 1A). HRT Atlas v1.0 offers a user friendly and reactive interface designed to provide a nice using experience. The web tool was tested in Firefox, Chrome, Explorer, Safari as well as android and iOS small devices.

## Database contents and features

### HRT Atlas v1.0 statistics

HRT Atlas analyzed 12,482 and 507 human and mouse samples respectively, involving 82 human cell and tissue types and 14 mouse cell and tissue types. After processing, 2,176 unique genes fulfilled our criteria and were defined as HK genes, of which 2,158 HK transcripts were constitutively expressed. In the mouse database, 3,277 HK transcripts and 3,024 HK genes fulfilled our criteria. At least two HK transcript-specific primers were designed per tissue to facilitate using of more than one candidate reference for normalization purpose. Overall, 44 and 23 primer-pairs were designed for human and mouse candidate reference genes, respectively.

Furthermore, 52% of the human HK genes present their orthologs in the mouse HK list (the full lists can be downloaded from Download page). We compared our list of HK genes with previously published lists which were also based on RNA-seq data sets (1, 25, 26). As shown in figure 2A, only 91 human HK genes were exclusively identified in the present study (Fig 2A). Similarly, a high overlap was also observed between previous mouse HK genes lists (45, 46) and our HRT Atlas database (Fig 2B). Importantly, thousands of genes identified in these previous studies didn’t fulfill our stringent criteria. These results support our hypothesis that these previous lists of HK genes, generated with low sample size and heterogeneous definitions of HK genes, might suffer from low specificity. We performed two simulations to investigate whether samples size and tissue type diversity can affect the prediction of HK genes/transcripts using GTEx dataset. Applying our HK transcripts detection criteria to 6 groups of random sampled tissues (5, 10, 15, 20, 30 and 50 tissue types respectively) we observed that the number of transcripts that fulfilled our HK transcripts criteria decreases as the number of tissue types increases (Supplementary figure 1A). With the second simulation, by using all of the tissues included in our analysis, we clearly observed that the accuracy of detection is also associated with the sample size (Supplementary figure 1B) as the stably expressed transcript’s number decreases with increasing sample size. For each of these simulations, random sample was obtained and 100 permutations were performed to detect the median number of stably expressed transcripts. Finally, we observed in figure 2A a large set of genes (1801 genes) that were detected by previous methods but not in HRT Atlas. This large genes set has been analyzed and we observed that about 98% didn’t fulfill the non-zero expression criterion. The remaining 2% have high variability that is a high standard deviation and/or MCF. The description of these genes in previous studies as housekeeping genes can result from the relatively small sample size of these studies, which as shown in the simulations, reduce the accuracy of the predictions. Together, these results and observation showed that both sample size and tissue/cell diversity can improve the accuracy of HK genes detection and explained at least partially the large discrepancy observed between previous studies and our database.

**Figure 2:**
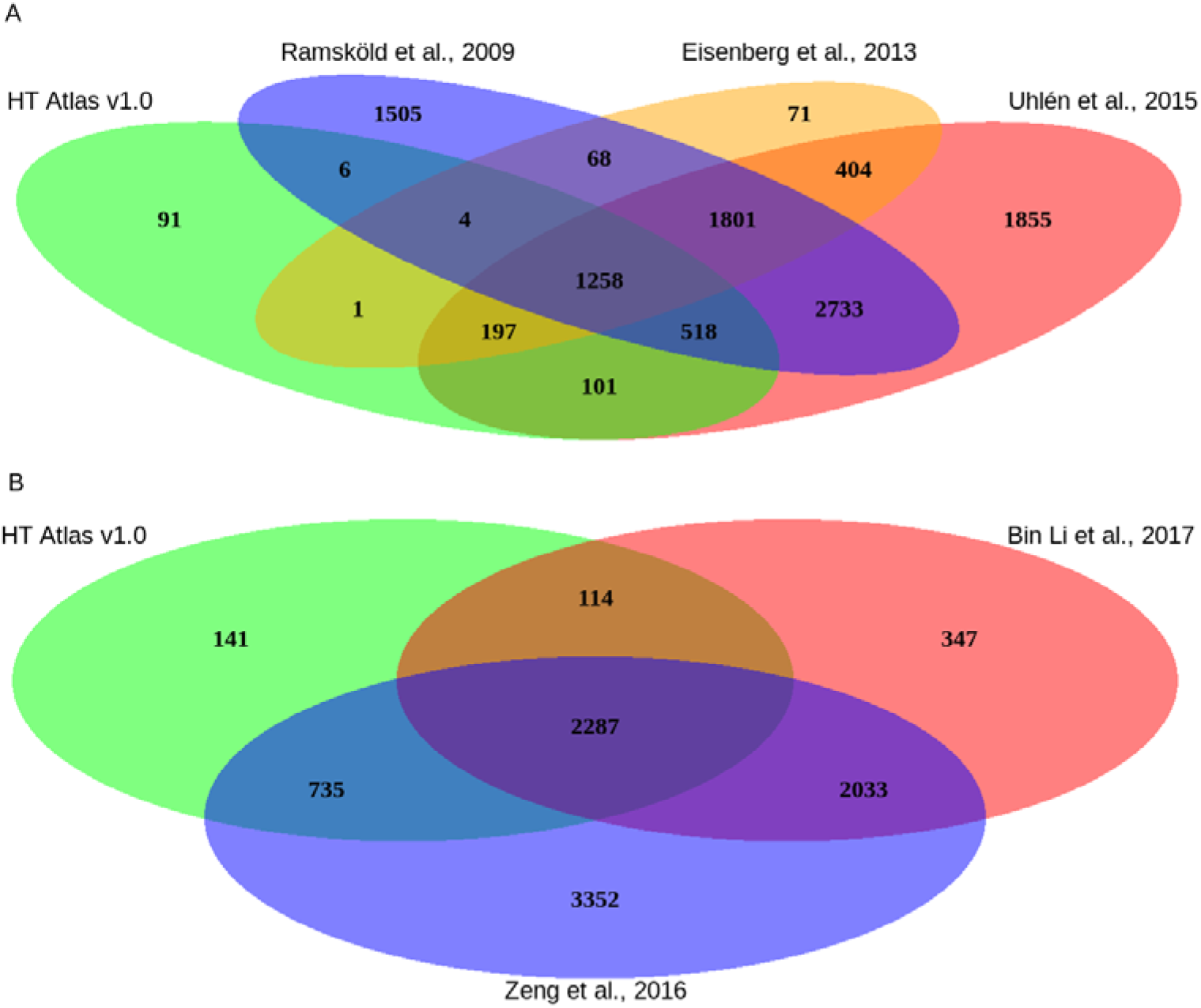
Overlap of HRT Atlas with other HK genes lists. (A) The diagram demonstrates that only 91 human genes were exclusively detected in the present study. In contrast, thousands of putative HK genes that have been previously listed by other authors (listed in the diagrams) were not identified by our criteria and are not listed in our HRT Atlas. (B) Similarly, when comparing HRT Atlas with previous mouse HK genes studies, 141 genes were exclusively detected in the present study, while several others which were listed in previous studies did not reach the criteria established in HRT Atlas.

### Web interface

From the home interface users can access human and mouse candidate reference genes databases by clicking on the human or mouse icons. The search interface allows searching for reliable candidate reference transcripts for qPCR normalization by selecting one of the 82 tissues or cell types included in our database. By clicking on the search box, the full list of cells and tissues is automatically displayed on the screen and user can select one option to interrogate the SQLite database. From the same interface users can also access link to visualize expression of HK transcripts across cells/tissues, or to download the complete list. A similar interface is provided for mouse database.

In each of these interfaces, after searching for a specific cell or tissue type, the results page shows the list of candidate reference transcripts. Each transcript is identified by its Ensembl ID, followed by the gene symbol. By default, results are ranked by the SP criteria described above. In addition, a color code is used to categorize transcripts according to expression levels (highly and moderately expressed in green and turquoise respectively). Alternatively, transcripts can be ranked at the convenience of the user by clicking in the arrow next to any of these parameters (Figure 3A). Users can select the number of candidate reference transcripts to be shown per page, and then download the full list displayed on the screen in their preferred format (csv, or pdf). Specific primer pair lists can also be downloaded. All the primer pairs were validated according to MIQE guidelines. User can access the quality parameters of each primer pair and download its standard curve and the melting curve.

**Figure 3:**
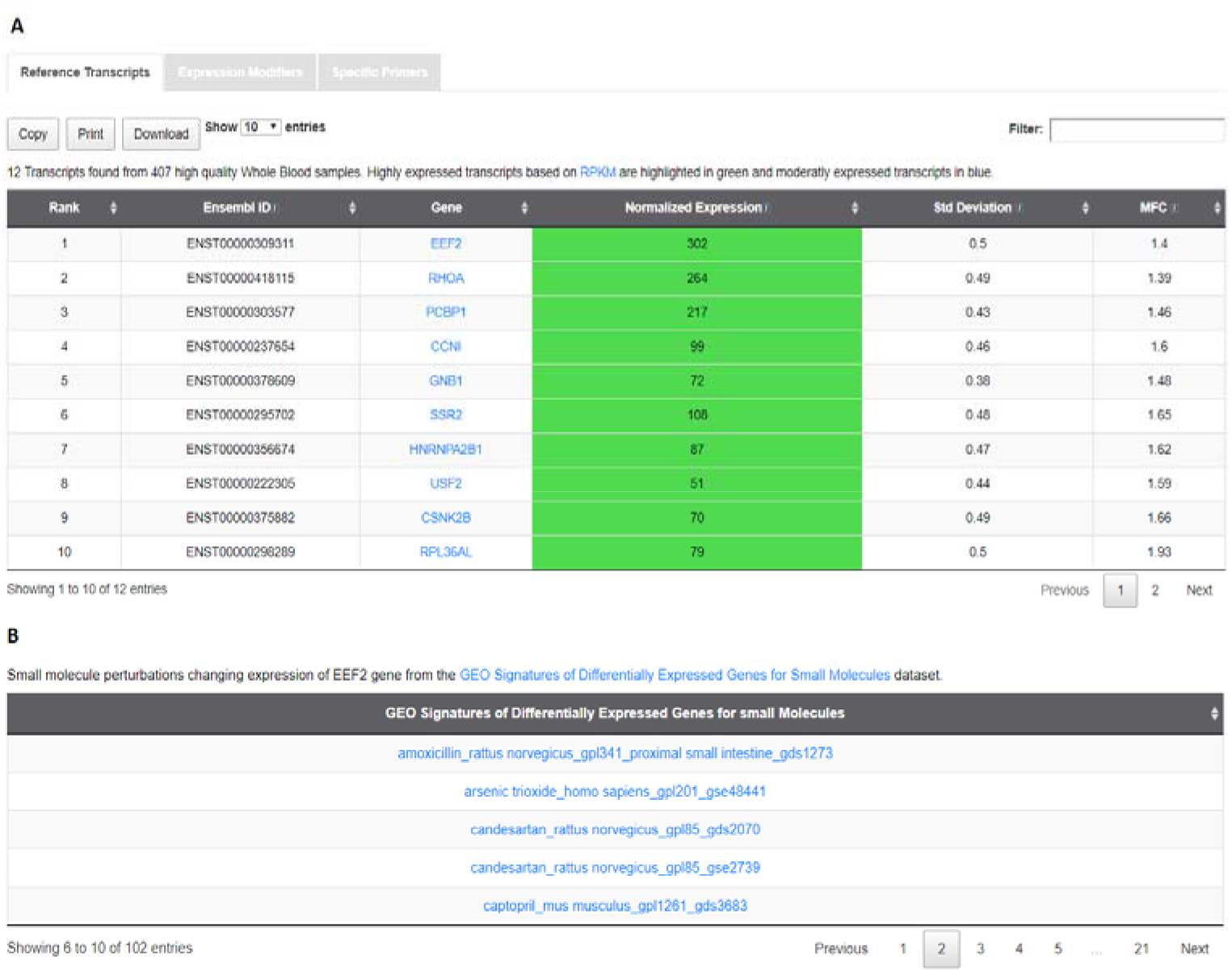
Screenshot of the candidate reference transcript search result page. (A) The result page presents a tab with three clickable menus (Reference Transcripts, Expression Modifiers, and Specific Primers). When clicking into this menu, the specified information is displayed. Panel A displays a list of 10 HK transcripts, ranked according to HRT Atlas algorithm. HK transcripts are identified by Ensembl ID and gene symbol. Quantitative parameters of gene expression are listed. (B) In the lower panel, a list of five potential modifiers (small molecules) of the expression of a specific HK gene is shown.

As diseases and small molecules are known to affect the stability of housekeeping genes, HRT Atlas is integrated with three manually curated functional association data sets from Harmonizome database that predict expression stability modifiers based on differential expression, and are intended to guide users in the choice of the best candidate reference genes for qPCR normalization. A gene must be avoided for normalization of qPCR if the disease or the molecule of interest appears in its modifiers list (Figure 3B). These datasets can be accessed from the results page. Alternatively, when clicking on “Expression Modifiers” in the navigation bar, users can access the same information for any single gene. In addition, HRT Atlas has been integrated with EpiRegio REST API (47) to retrieve information about regulatory elements (REMs) that are able to regulate gene expression. This resource can also be accessed from results page (Regulatory Element). EpiRegio is a web server which allows the analysis of genes and their associated REMs estimated in cell type-specific context.

Users can access the “Gene expression” menu from the navigation bar to visualize gene expression across tissue and cell types. This interface also provides the description of a selected HK gene, its official name and its synonyms retrieved from the NCBI’s portal and based on RefSeq project. When a gene was identified as a reliable candidate reference gene for a subset of tissues and cells, a list of that tissues and cell types is also provided (Fig 4).

**Figure 4:**
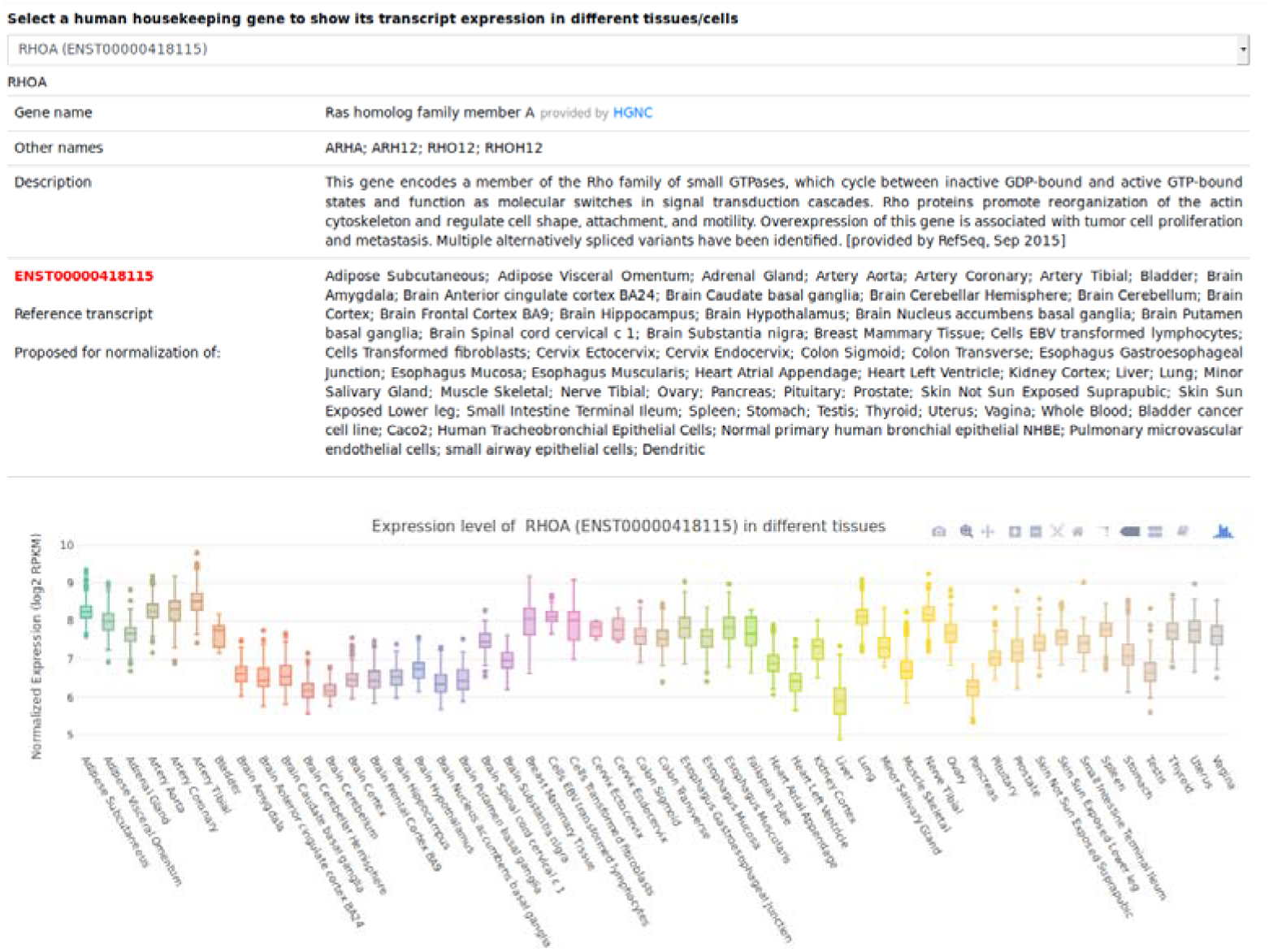
Screenshot of the expression visualization page. For each selected HK gene, information such as gene nomenclature, synonyms and a list of tissues with stable expression are listed. Box plots depicting transcript expression in different cells and tissues using data from our database are also shown. Boxplots were generated using plotly package (Carson Sievert (2018) plotly R. https://plotly-r.com).

Finally, by clicking on the “Download” menu from the navigation bar, users can download the complete list of human and mouse HK genes and HK transcripts. From the same page users can download a list of the most stable genes across all non-disease analyzed tissues. These genes can be validated in cell types not included in our database for further normalization purposes.

### Case study of the use of the database

The main application of the database is qPCR gene expression normalization. As a proof of concept of the reliability of the newly proposed candidate reference transcripts, two candidate reference transcripts described in PBMCs (the transcripts ENST00000253039 and ENST00000335508 for EIF2S3 and SF3B1) were tested by comparing their stability metrics (geNorm M metric) with some commonly used genes (ACTB, B2M, GAPDH and PPIA1). Our results (geNorm ranking) showed that the newly described transcripts in PBMCs were the most suitable for normalization in this testing context (Figure 5). Interestingly GAPDH, in 3th position, has been described as HK transcript in the database but not as a suitable candidate reference gene in PBMC (Figure 5). This observation reinforces the necessity of tissue-selective section. Alternatively, we also used RNAseq datasets to calculate the same stability metrics across 12 different tissues random sampled that were included in our database (Supplementary Table 1). This analysis compared the best ranked candidate reference transcripts in each tissue with: (i) their respective genes (included all of their coding and non-protein coding transcripts) and (ii) 12 other commonly used reference genes. As shown in the supplementary table 1, the newly described candidate reference transcripts were more suitable for normalization than genes commonly used. Interestingly, the candidate reference transcripts also exhibited greatest stability in comparison with their respective genes expression. These results also suggested that the reference transcript-based normalization strategy instead of the commonly used gene-based method and the newly proposed candidate references transcript may be considered as suitable method of gene expression normalization.

**Figure 5:**
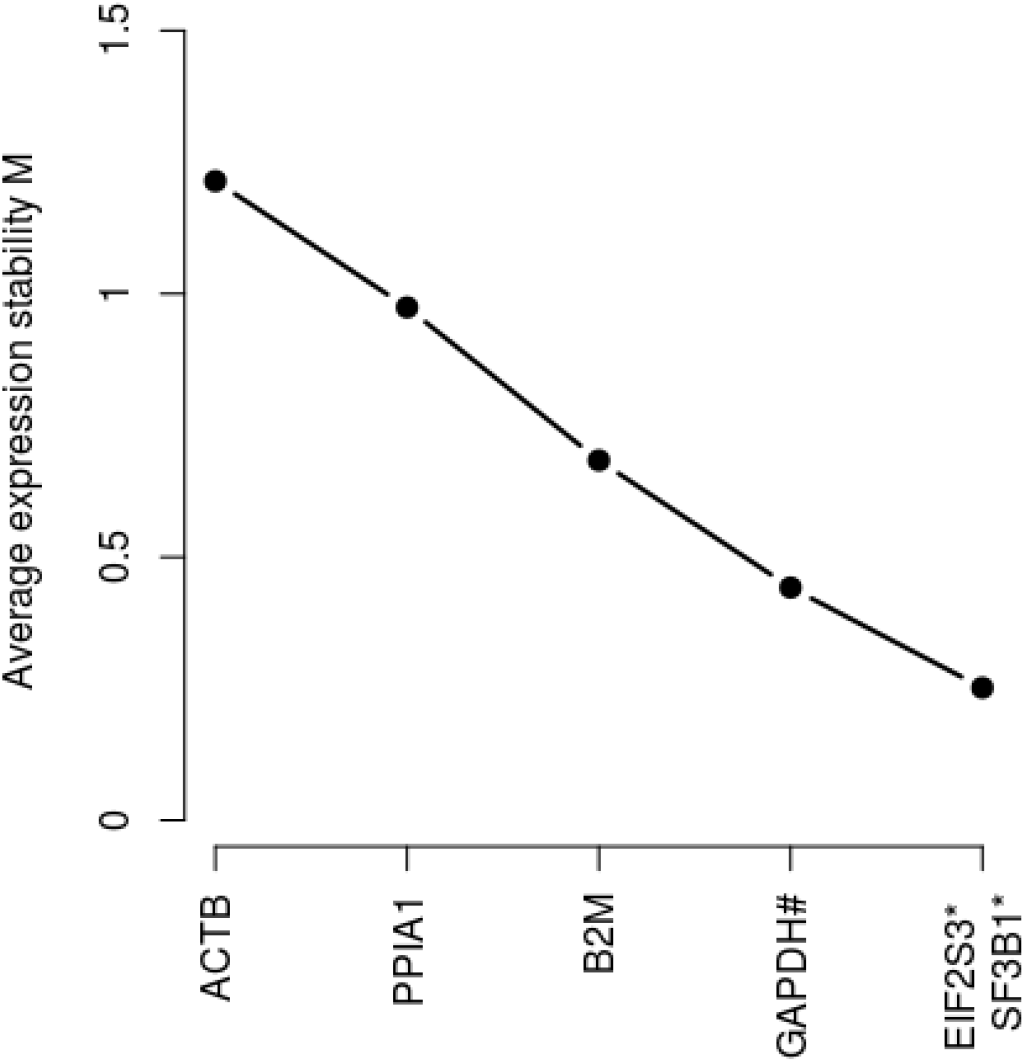
Two candidate reference transcripts proposed for PBMCs were most suitable for normalization in comparison with commonly used reference genes. The performance of newly described candidate reference transcripts and proposed for PBMCs (the transcripts ENST00000253039 and ENST00000335508 for EIF2S3 and SF3B1) were compared with some commonly used genes (ACTB, B2M, GAPDH and PPIA1). The combination of transcripts ENST00000253039 and ENST00000335508 is the most suitable with the lowest geNorm M stability metric. 7 samples were analyzed. *: gene names of the new reference transcripts.

HRT Atlas database has a limitation that need to be acknowledged. The relative small sample size of mouse datasets can reduce the accuracy of the prediction of mouse housekeeping or reference transcripts. Unfortunately, there are no large datasets with homogeneously pre-processing for mouse as the GTEx datasets for humans so that RNA-seq data have been manual curated. As we analyzed expression at transcript level, we opted to include only datasets that fulfilled some criteria known to contribute to a more robust transcript’s expression estimate such as: (i) the inclusion of libraries only constructed with paired-end read to improve estimation accuracy over single-end reads (48, 49); (ii) high sequencing depth with at least twenty millions of reads to improve the estimation of low abundant genes and exons and splice junctions (34). Furthermore, only wild type mice were included. So, by prioritizing these quality criteria over a large but heterogeneous dataset, we aimed to minimize unwanted experimental confounder in our predictions. In our knowledge, this is the largest mouse dataset analyzed in a workflow of mouse HK genes detection. Despite this small data size, human and mouse databases have 52% of detected orthologous genes. Furthermore, the HK genes described in human as well as in mouse were shown to be involved mainly in basal metabolism pathways (Supplementary table 2 and 3) such that one can predict that they are enriched in HK genes. Finally, the prediction of gene modifiers will be improved in future versions. Because there is no way to know whether gene expression will change under all experimental and disease conditions, we recommend empirical validation of the proposed candidate reference transcripts before using in qPCR experiments. Furthermore, we highly recommended users to consider using of at least two candidate reference transcripts for qPCR normalization as recommended by MIQE guidelines.

In conclusion, HRT Atlas v1.0 represents a valuable tool for researchers from a wide range of fields in biomedical research, due to its capability to refine the identification of a critical parameter (i.e. the gene used to calibrate expression level reads) in one of the most commonly used techniques of molecular biology studies. The database can also be used to assist in research questions about structural and functional genomics that require a more precise identification of human and mouse HK genes and transcripts. Our strategy for the future will focus in including more cells and tissues into the database. We are also planning to analyze, using the same workflow, samples from other model organisms.

## Acknowledgments

The Genotype-Tissue Expression (GTEx) Project was supported by the Common Fund of the Office of the Director of the National Institutes of Health, and by NCI, NHGRI, NHLBI, NIDA, NIMH, and NINDS. Part of the data used for the analyses described in this manuscript was obtained from GTEx v7.

## Data availability

All data are available from HRT Atlas (http://www.housekeeping.unicamp.br). Processing codes and source codes are available from github (https://github.com/Bidossessih/housekeepingAtlas).

