## Supplemental Figure 1 for "HRT Atlas v1.1 database: redefining human and mouse housekeeping genes and candidate reference transcripts by mining massive RNA-seq datasets"

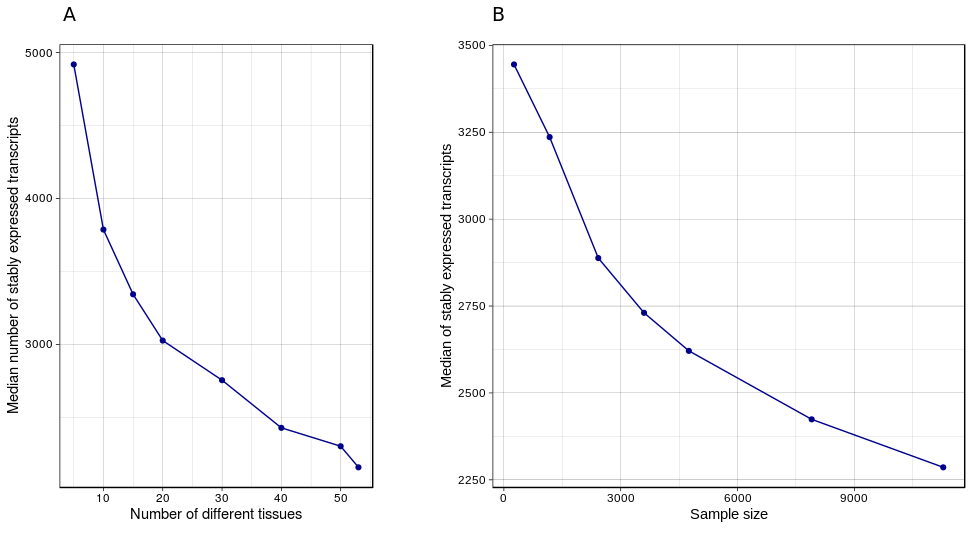

Figure 1: Simulation of the impact of tissue diversity and samples size.

Simulations using GTEx dataset based on random sampling and 100 permutations have been performed to detect the median number of stably expressed transcripts. Our results showed that tissue type diversity (A) and samples size (B) can affect the prediction of HK genes/transcripts. In both simulations the number of transcripts that fulfilled HRT Atlas criteria decreases as the number of tissue types (A) or sample size increases (B).
