## Supplemental Table 1 for "HRT Atlas v1.1 database: redefining human and mouse housekeeping genes and candidate reference transcripts by mining massive RNA-seq datasets"

**HRT Atlas v1.1 database: redefining human and mouse housekeeping genes and candidate reference transcripts by mining massive RNA-seq datasets**
Bidossessi Wilfried Hounkpe1, Francine Chenou1, Franciele Lima1, Erich Vinicius de Paula1,2
Affiliations: 1 School of Medical Sciences, University of Campinas, Campinas, SP, Brazil; 2 Hematology and Hemotherapy Center, University of Campinas, Campinas, SP, Brazil

**GeNorm stability metrics across 12 different tissues based in RNA-seq datasets**

**Legend**

Gene level expression of the selected reference transcripts (included all of their coding and non-protein coding transcripts)

Newly described reference transcripts

Commonly used reference genes

| **Rank** | **Artery Aorta** | **Brain Cerebellar Hemisphere** | **Brain Cerebellum** |
| --- | --- | --- | --- |
| 1 | PCBP1 | ENST00000358704 | ENST00000316509 |
| 1 | ENST00000382581 | ZBTB18 | ENST00000447750 |
| 3 | ENST00000322535 | ENST00000378609 | ENST00000378609 |
| 4 | ENST00000447750 | ENST00000300086 | ENST00000309311 |
| 5 | ENST00000292807 | ENST00000316509 | ENST00000375882 |
| 6 | ENST00000175091 | ENST00000447750 | ENST00000334478 |
| 7 | ENST00000418115 | ENST00000309311 | ENST00000257013 |
| 8 | ENST00000332556 | ENST00000356674 | ENST00000360472 |
| 9 | ENST00000309311 | ENST00000373232 | ENST00000356674 |
| 10 | ENST00000216181 | ENST00000250894 | HPRT1 |
| 11 | LAPTM4A | TERF2IP | RTL8C |
| 12 | LAMP1 | PPA1 | TBP |
| 13 | HPRT1 | PGK1 | PGK1 |
| 14 | TBP | HPRT1 | CSNK2B |
| 15 | MRFAP1 | TBP | VAMP2 |
| 16 | RHOA | VAMP2 | PEA15 |
| 17 | GUSB | HNRNPA2B1 | GNB1 |
| 18 | PPIA | PPIA | HNRNPA2B1 |
| 19 | PGK1 | GUSB | GUSB |
| 20 | SF3B2 | ACTB | PPIA |
| 21 | RPS18 | RPS18 | RPS18 |
| 22 | GDI1 | EEF2 | GDI1 |
| 23 | EEF2 | GAPDH | ACTB |
| 24 | MYH9 | TFRC | TFRC |
| 25 | AP2M1 | GNB1 | GAPDH |
| 26 | YWHAZ | GDI1 | PFDN5 |
| 27 | GAPDH | RPLP0 | EEF2 |
| 28 | ACTB | B2M | RPLP0 |
| 29 | TFRC | MAPK8IP3 | YWHAZ |
| 30 | B2M | YWHAZ | B2M |
| 31 | RPLP0 |  |  |

| **Rank** | **Brain Cortex** | **Breast Mammary Tissue** | **Cervix Ectocervix** |
| --- | --- | --- | --- |
| 1 | ENST00000249289 | ENST00000322535 | PCBP1 |
| 1 | ENST00000357156 | ENST00000392132 | ENST00000418115 |
| 3 | ENST00000257013 | ENST00000382581 | ENST00000357156 |
| 4 | ENST00000447750 | ENST00000502976 | ENST00000309311 |
| 5 | ENST00000382581 | ENST00000175091 | ENST00000409614 |
| 6 | ENST00000037243 | ENST00000330720 | ENST00000376630 |
| 7 | ENST00000356674 | ENST00000265062 | ENST00000412585 |
| 8 | ENST00000577035 | ENST00000311481 | ENST00000376809 |
| 9 | ENST00000360472 | ENST00000330938 | DYNLRB1 |
| 10 | ENST00000371646 | RAB1B | ENST00000368719 |
| 11 | ATP6V1F | ENST00000356674 | ENST00000371646 |
| 12 | DYNLRB1 | HPRT1 | HPRT1 |
| 13 | MRFAP1 | LAPTM4A | HSP90AB1 |
| 14 | HPRT1 | MRFAP1 | RHOA |
| 15 | TBP | TBP | TBP |
| 16 | GABARAPL2 | XRCC5 | HLA-E |
| 17 | RTL8C | KDELR1 | HLA-A |
| 18 | PGK1 | PTTG1IP | GUSB |
| 19 | PEA15 | HNRNPA2B1 | TFRC |
| 20 | HSP90AB1 | SF3B2 | ACTB |
| 21 | GABARAP | PGK1 | HLA-B |
| 22 | HNRNPA2B1 | RAB7A | EEF2 |
| 23 | GUSB | CNBP | SERF2 |
| 24 | PPIA | GUSB | B2M |
| 25 | RPS18 | PPIA | S100A6 |
| 26 | GDI1 | RPS18 | PGK1 |
| 27 | GAPDH | TFRC | PPIA |
| 28 | TFRC | YWHAZ | RPS18 |
| 29 | ACTB | GAPDH | RPLP0 |
| 30 | RPLP0 | B2M | GAPDH |
| 31 | YWHAZ | ACTB | YWHAZ |
| 32 | B2M | RPLP0 |  |

| **Rank** | **HEK293** | **PANC1** | **RKO** |
| --- | --- | --- | --- |
| 1 | GHITM | ENST00000361611 | ENST00000229563 |
| 1 | ENST00000373232 | ENST00000451728 | ENST00000361611 |
| 3 | ENST00000262193 | ENST00000229563 | ENST00000315436 |
| 4 | ENST00000524896 | HPRT1 | ENST00000503362 |
| 5 | ENST00000373365 | TBP | ENST00000340941 |
| 6 | GLO1 | GUSB | ENST00000451728 |
| 7 | PSMB1 | TMEM14C | ENST00000290649 |
| 8 | ENST00000262225 | PSMB5 | TBP |
| 9 | ENST00000253039 | CNBP | TMEM14C |
| 10 | ENST00000328848 | PPIA | GUSB |
| 11 | ENST00000321301 | ACTB | PPIA |
| 12 | NOP10 | PGK1 | AMFR |
| 13 | ENST00000299767 | TFRC | PSMB5 |
| 14 | PPA1 | GAPDH | TFRC |
| 15 | EIF2S3 | YWHAZ | SPCS3 |
| 16 | TMED2 | B2M | WSB2 |
| 17 | HPRT1 | RPLP0 | GAPDH |
| 18 | TBP | RPS18 | HPRT1 |
| 19 | TOMM5 |  | PGK1 |
| 20 | TFRC |  | MCCC2 |
| 21 | PGK1 |  | RPLP0 |
| 22 | GUSB |  | ACTB |
| 23 | HSP90B1 |  | CNBP |
| 24 | B2M |  | B2M |
| 25 | PPIA |  | YWHAZ |
| 26 | EIF3M |  | RPS18 |
| 27 | ACTB |  |  |
| 28 | GAPDH |  |  |
| 29 | RPLP0 |  |  |
| 30 | YWHAZ |  |  |
| 31 | RPS18 |  |  |

| **Rank** | **SKBR3** | **Stomach** | **Whole Blood** |
| --- | --- | --- | --- |
| 1 | ENST00000229563 | ENST00000265062 | PCBP1 |
| 1 | ENST00000322428 | ENST00000332556 | ENST00000378609 |
| 3 | ENST00000361611 | ENST00000435120 | ENST00000418115 |
| 4 | ENST00000340941 | ENST00000328024 | ENST00000222305 |
| 5 | ENST00000256255 | ENST00000315758 | ENST00000375882 |
| 6 | ENST00000395068 | LAMP1 | RPL36AL |
| 7 | TBP | ENST00000330938 | ENST00000295702 |
| 8 | TMEM14C | ENST00000237530 | ENST00000237654 |
| 9 | MAF1 | ENST00000301740 | ENST00000356674 |
| 10 | GUSB | HPRT1 | ENST00000309311 |
| 11 | MCCC2 | RAB11B | HPRT1 |
| 12 | HPRT1 | TBP | TBP |
| 13 | PGK1 | PGK1 | CCNI |
| 14 | SARAF | RAB7A | RHOA |
| 15 | TFRC | MDH2 | CSNK2B |
| 16 | RPLP0 | RPN2 | HNRNPA2B1 |
| 17 | PSMB5 | MLF2 | PGK1 |
| 18 | PPIA | PPIA | GUSB |
| 19 | GRINA | PTTG1IP | GNB1 |
| 20 | GAPDH | RPS18 | SSR2 |
| 21 | B2M | GUSB | USF2 |
| 22 | ACTB | TFRC | EEF2 |
| 23 | YWHAZ | YWHAZ | PPIA |
| 24 | RPS18 | GAPDH | RPS18 |
| 25 |  | RPLP0 | YWHAZ |
| 26 |  | B2M | TFRC |
| 27 |  | SRRM2 | ACTB |
| 28 |  | ACTB | B2M |
| 29 |  |  | GAPDH |
| 30 |  |  | RPLP0 |
